# Mapping the habitat suitability of *Culex pipiens* in Europe using ensemble bioclimatic modelling

**DOI:** 10.1101/2025.03.17.643724

**Authors:** Lara Marcolin, Agnese Zardini, Eleonora Longo, Beniamino Caputo, Piero Poletti, Moreno Di Marco

## Abstract

Anthropogenic pressure on natural ecosystems has profoundly influenced the dynamics of disease vectors, altering their distribution and phenology, and increased the risk of vector-borne diseases. *Culex pipiens* is a widespread mosquito species and an important vector of zoonotic pathogens in Europe, including West Nile virus. Understanding *Cx. pipiens* distribution is essential to orient monitoring and mitigation strategies for vector-borne disease risk. Here, we employed an ensemble modelling framework to investigate the climatic and environmental determinants of *Cx. pipiens* distribution in continental Europe. We integrated high-resolution occurrence data from entomological surveillance with bioclimatic, topographic, and anthropogenic variables. Our findings identified imperviousness as the most influential predictor of *Cx. pipiens* distribution, with a strong association with human-modified low-elevation areas that reflects its adaptability to anthropogenic environments. The ensemble prediction approach we employed demonstrated higher predictive accuracy and transferability compared to individual models during independent block validation. These results emphasize the need to incorporate anthropogenic factors into disease vector distribution models to support evidence-based surveillance and control strategies, while also offering updated, robust, and spatially explicit predictions of the habitat suitability for *Cx. pipiens* in Europe. Overall, this study highlights the role of human modification of the natural environment in shaping *Cx. pipiens* distribution, extending previous knowledge on the role of urban areas.

## 1. Introduction

*Culex pipiens* is a widely distributed mosquito native to Africa, Asia, and Europe, and found across temperate regions worldwide, including North and South America, Australia, and the Middle East (Harbach, 2012). In Europe, *Cx. pipiens* is found in all countries except Iceland and Faroe Island (ECDC, 2023). This species thrives in both rural and urban environments, adapting to diverse habitats that range from rural to highly urbanized landscapes (Krol et al., 2023). Since *Cx. pipiens* feeds on a broad range of vertebrate hosts, including birds, humans, and other mammals, it is an important vector of several zoonotic pathogens. In Europe, these include West Nile, Usutu, and Sindbis viruses (Turell, 2012), along with canine dirofilarial worms, such as *Dirofilaria immitis* and *D. repens* (Bravo-Barriga et al., 2016; Younes et al., 2021). Moreover, *Cx. pipiens* mosquitoes have been identified as moderately to highly competent vectors of Rift Valley fever and Japanese encephalitis virus under laboratory conditions (De Wispelaere et al., 2017; Vloet et al., 2017) and should be considered as potential vectors of these viruses in Europe. Given the prominent role in the transmission of several diseases, *Cx. pipiens* is a critical species for public health surveillance in Europe. Understanding the ecological and environmental variables driving its distribution is therefore important for anticipating and mitigating vector-borne disease risk. However, this requires careful consideration of species biology for informing input variable selection, as including non-relevant variables in species distribution modelling can hamper the consistency and reliability of predictions (Santini et al., 2021).

By combining species occurrence data with environmental and climatic variables, species distribution models (SDMs) elucidate the ecological drivers underpinning species distributions and offer spatially explicit predictions of habitat suitability (Elith & Leathwick, 2009). These tools can help identify areas at risk of vector-borne disease transmission, to inform targeted surveillance and control strategies. The application of SDMs to vectors like *Cx. pipiens* is essential for understanding how anthropogenic and environmental factors interact to shape their distributions and to anticipate changes under future scenarios.

Previous studies have attempted to model the spatial distribution of *Cx. pipiens* in Europe (Versteirt et al., 2013; Wint et al., 2020), but they either did not employ an ensemble approach or, if they did, they did not explicitly validate the ensemble predictions. Specifically, these studies often lack robust evaluation, e.g. in the form of spatial block cross-validation or independent new-label datasets. Moreover, they do not optimize the binarization threshold for the classification of areas as suitable or unsuitable, relying instead on arbitrary and suboptimal threshold, which can affect model accuracy and interpretability. These limitations are critical, as SDMs are prone to overfitting, leading to overly optimistic predictions. To mitigate this risk, model validation must be independent of training data.

Ensemble modelling provides a powerful framework to address these challenges, as it reduces dependency on any single algorithm and accounts for the variability among different modelling techniques (Thuiller et al., 2009, 2019). By aggregating multiple predictions, ensemble approaches enhance model robustness, limit overfitting, and yield more generalizable results. Moreover, they improve predictive accuracy by balancing different sources of bias and uncertainty, making them particularly well suited for ecological applications (Elith & Leathwick, 2009).

In this work, we investigate the environmental determinants of the distribution of *Cx. pipiens* in Europe. To this end, we implement an ensemble modelling framework that integrates multiple techniques to reduce methodological uncertainty and generate reliable predictions. We validate our model ensemble using both new-label validation and spatial block cross-validation. Additionally, we optimize the binarization threshold, ensuring a more robust and accurate model.

## 2. Methods

### 2.1. Data collection

#### 2.1.1. *Cx. pipiens* occurrence and background data

We retrieved occurrence data for *Cx. pipiens* in Europe from the VectorNet project archive (Wint et al., 2023), a joint initiative of entomological surveillance commissioned by the European Centre for Disease Prevention and Control (ECDC) and the European Food Safety Authority (EFSA). This archive provides a comprehensive dataset of records from entomological traps targeting key vectors of pathogens relevant to human and animal health in Europe, such as ticks, mosquitoes, sand flies, and biting midges. For this study, the dataset was filtered to include only confirmed occurrences of *Cx. pipiens* recorded between 2008 and 2018. Each occurrence point represents a trap location with precise GPS coordinates where at least one adult individual of *Cx. pipiens* was successfully captured during the study period. The initial dataset retrieved from the VectorNet archive contained n=1105 presence points of *Cx. pipiens* across Europe. As the data were affected by spatial clustering patterns (i.e. associated to monitoring efforts in certain localities), we filtered the dataset by retaining a single (randomly selected) occurrence point per each 10x10km area, thereby ensuring a more even spatial distribution of presence points. We then defined background points by randomly selecting points within biogeographic ecoregions (Olson et al., 2001) containing at least one confirmed occurrence point after excluding areas within a 1 km buffer around each presence point (Whitford et al., 2024), resulting in a total of 497 presence and 4970 absence points.

#### 2.1.2. Environmental covariates

We selected a set of bioclimatic and environmental covariates associated with *Cx. pipiens* occurrence, based on the ecological requirements of the species (Gangoso et al., 2020; Gorris et al., 2021). Selected variables included an elevation map for Europe (Fick & Hijmans, 2017), climatic metrics such as relative humidity, climate moisture index, potential evapotranspiration, and bioclimatic variables derived from CHELSA database (Karger et al., 2017) with a spatial resolution of 30 arcseconds. The selected bioclimatic variables from CHELSA included annual mean temperature (BIO01), temperature seasonality (BIO04), mean temperature of the wettest quarter (BIO08), mean temperature of the driest quarter (BIO09), annual precipitation (BIO12), precipitation seasonality (BIO15), precipitation of the wettest quarter (BIO16), and precipitation of the driest quarter (BIO17). These variables were chosen for their direct influence in modelling *Cx. pipiens* habitat suitability, particularly with regard to the thermal and hydrological conditions favourable for *Cx. pipiens* breeding and development (Gangoso et al., 2020; Gorris et al., 2021). To further refine the models, land cover and human population density variables were incorporated to provide a nuanced representation of habitat and socio-environmental factors driving the spatial distribution of *Cx. pipiens* (Amdouni et al., 2022; Krol et al., 2023).

Monthly data on precipitation, minimum temperature, and maximum temperature were aggregated for each year between 2008 and 2018 and subsequently used to derive bioclimatic variables using the “dismo” package in R (Hijmans et al., 2023). These annual values were then averaged over the entire study period (2008–2018) to obtain a single mean value for each variable. Similarly, monthly data on relative humidity, climate moisture index, and potential evapotranspiration were aggregated into annual values and averaged across the study period. Given its importance as a predictor of *Cx. pipiens* distribution at country scale in Europe (Gangoso et al., 2020), imperviousness for the 2012 reference year was derived from the Copernicus Land Monitoring Service high-resolution layers (https://land.copernicus.eu/pan-european) and included as a proxy for human infrastructure (including buildings, roads, etc.) which represent an impermeable barrier between the soil and above-ground environment, preventing rainwater from penetrating the ground. This map quantifies sealing density in the range from 0% to 100% at a spatial resolution of 100 m. Cropland land cover data for the 2015 reference year were also derived from the Copernicus Land Monitoring Service high-resolution layers (https://land.copernicus.eu/pan-european) at a spatial resolution of 100 m. Human population data with a spatial resolution of 30 arcseconds were obtained from the Gridded Population of the World, Version 4 (GPWv4; CIENSIN, 2017). This dataset provides high-resolution estimates of population density (persons per square kilometer). All variables were aggregated to a 1 km resolution for analysis.

To prevent multicollinearity among predictors, variance inflation factor (VIF) analysis (Naimi et al., 2014) was performed through a stepwise procedure, retaining only variables with VIF < 3 for subsequent modelling.

### 2.2. Species distribution models

We applied an ensemble modelling approach using the “biomod2” package (Thuiller et al., 2024) in R, a framework that enables robust predictions of species distributions by integrating multiple modelling techniques to address methodological uncertainties and explore species-environment relationships. We implemented five distinct models: Generalized Additive Models (GAM), Neural Network (ANN), Random Forest (RF), Maxnet, and Extreme Gradient Boosting (XGBOOST). Model-specific parameters were fine-tuned (Table S1) by testing a range of potential values and selecting the configuration that maximized the Boyce index (Boyce et al., 2002).

We evaluated model reliability and spatial transferability using a spatial block cross-validation, implemented through a dedicated function within the “biomod2” package. This method partitions both presence and background points into four spatial bins by dividing the study area along latitude and longitude, ensuring an even distribution of occurrence localities across the partitions (Muscarella et al., 2014). For the ensemble prediction, we retained only models achieving a Boyce index of 0.5 or higher (on a -1 to +1 scale). The final ensemble output was generated as a weighted average of the predictions of each individual model, with weights proportional to each model’s Boyce index (in order to prioritize higher-performing predictions).

To further validate the ensemble prediction, we performed a second round of spatial block cross-validation, this time applied to the final ensemble output rather than the individual models. Specifically, we used the same data partitions initially employed to validate each separate model, but here we assessed the predictive performance of the aggregated ensemble predictions. Additionally, we conducted a “new-label” validation conducted using independent occurrences of *Cx. pipiens* not used to train our model. We extracted new label records from the GBIF database, focusing on records from 2008 to 2018, corresponding to our study period (GBIF, 2024). The dataset was first filtered to retain only records with valid geographic coordinates. Occurrences were subjected to a series of quality control tests using the “CoordinateCleaner” package in R (Zizka, 2017). These tests identified potential issues, including records in capital cities, centroids of administrative regions, open ocean areas, the GBIF headquarters, urban areas or occurrences associated to biodiversity institutions. Additionally, these tests flagged zero coordinates, identical latitude or longitude and invalid coordinates, duplicates, or extreme outliers. Problematic records were excluded based on the outcome of these tests, resulting in a final dataset of n=436 *Cx. pipiens* occurrences in Europe.

By default, the “biomod2” package calculates performance metrics for each individual model by binarizing continuous probability values based on the threshold leading to the best True Skill Statistic (TSS; Allouche et al., 2006) score. Similarly, for the block validation and new-label validation of ensemble prediction, as well as for the final ensemble map, we selected the binarization threshold that maximized the TSS of the model ensemble, ensuring optimal discrimination between suitable and unsuitable areas. We estimated variable contribution to each model’s prediction based on permutation importance. The influence of each variable on model’s predictions was assessed by randomly permuting its value and measuring the correlation between original and shuffled variables (lower correlation indicating higher variable importance). To ensure robust estimates, the permutation process was repeated five times. We also generated response curves to explore the relationship between the covariates and the predicted probability of presence of *Cx. pipiens* (Elith et al., 2005).

Based on the ensemble model predictions, we generated a map displaying the continuous probability of *Cx. pipiens* presence and a binary habitat suitability map, obtained by applying the optimized threshold to the predicted probabilities. To ensure a realistic spatial representation, we limited both maps to the latitudinal range encompassing 99% of known *Cx. pipiens* occurrences recorded in GBIF (GBIF, 2024) since 1950, and in VectorNet (Wint et al., 2023) since 1980.

## 3. Results

### 3.1. Model performance

After applying the variance inflation factor (VIF) analysis to address multicollinearity among predictors, we retained nine environmental variables with VIF values < 3 (Table S2). These variables included temperature seasonality (BIO04, VIF = 1.49), mean temperature of the wettest quarter (BIO08, VIF = 2.07), precipitation seasonality (BIO15, VIF = 1.76), precipitation of the wettest quarter (BIO16, VIF = 1.84), elevation (VIF = 1.95), relative humidity (VIF = 1.91), imperviousness (VIF = 1.64), human population (VIF = 1.55), and cropland (VIF = 1.39).

We retained all models for ensemble predictions as they met the minimum performance threshold of Boyce index ≥ 0.5 during cross-validation (Figure 1a, Table S3).

**Figure 1.**
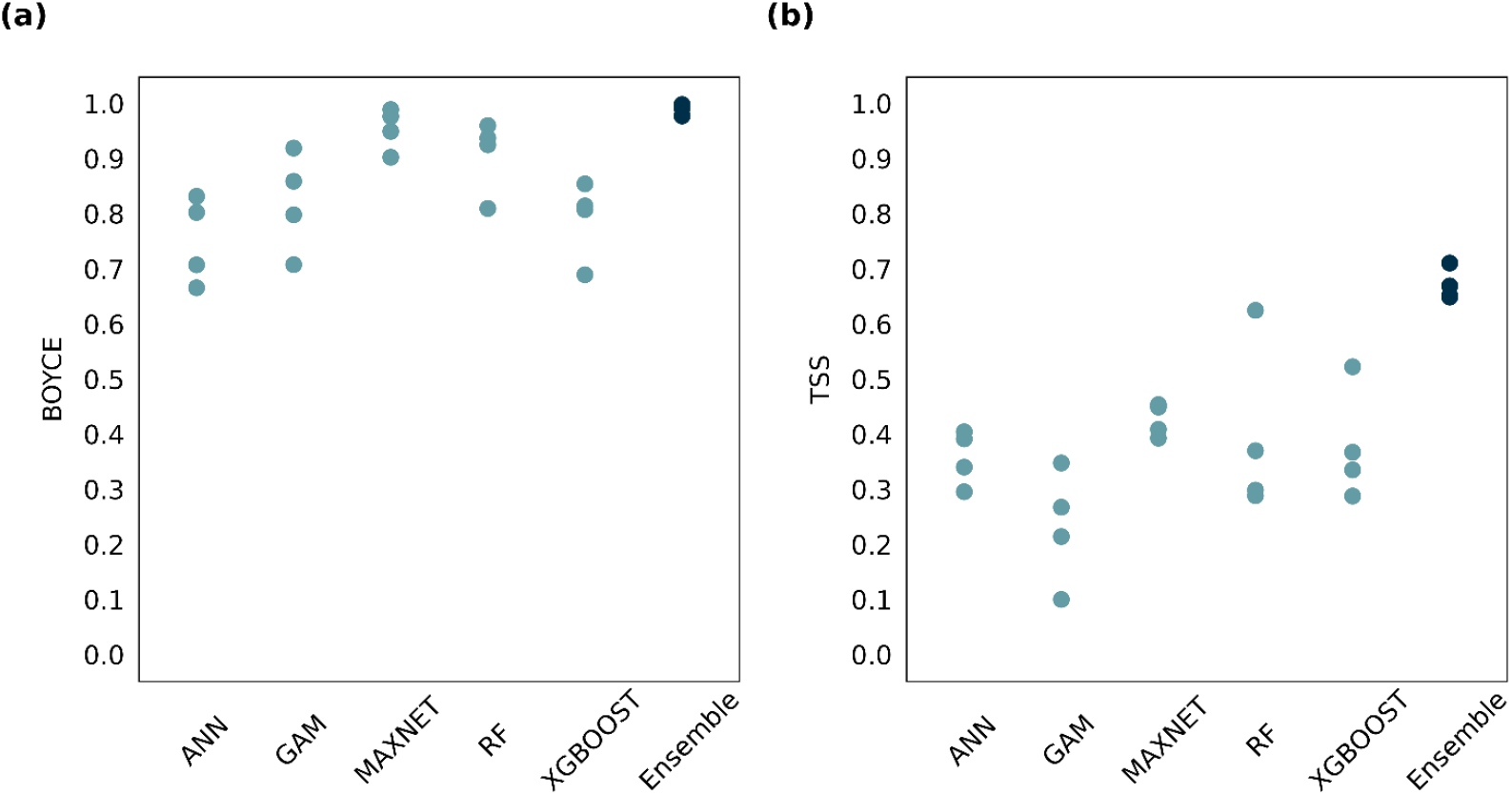
Model validation. Boyce index (a) and TSS (b) scores for each algorithm included in the ensemble model, as well as for the final ensemble. For the ensemble prediction, we retained only runs achieving a Boyce index of 0.5 or higher. Dots represent Boyce index and TSS scores (on a -1 to +1 scale) for individual runs calculated on the spatial block cross-validation data.

The final ensemble predictions were validated through a spatial block cross-validation and a new-label validation, after selecting 0.37 as the optimal probability threshold to classify suitable from unsuitable areas (leading to the highest TSS of 0.67; Figure S1a). Notably, in the block validation, the threshold of 0.37 led to much closer values of correctly predicted presences and correctly predicted absences (respectively 0.92 and 0.80) compared to a default binarization threshold of 0.5. We found higher predictive performance of the ensemble model compared to individual models (Figure 1b), both when using the optimal threshold to binarize ensemble predictions and when using the default 0.5 threshold (TSS = 0.67 vs TSS = 0.62; Figure S1b). In addition, the ensemble also showed a higher mean Boyce index of 0.991 (SD = 0.009) when compared to the mean values of individual models, which ranged from 0.76 for ANN to 0.96 for RF.

In the “new-label” validation, the median probability of presence was higher for independent GBIF occurrence points compared to randomly generated background points (Figure S2). This difference was statistically significant, as confirmed by a two-sample Kolmogorov-Smirnov test (D = 0.41392, p < 2.2 × 10^−16^). The test indicated a substantial divergence in the distribution of suitability values between occurrence and background points, suggesting that the model effectively distinguished presence locations.

### 3.2 Variable importance and response curves

The contribution of environmental variables to the predictions varied across the four ensemble runs, as reflected in the weighted permutation importance scores (Figure 2, Figure S3). Imperviousness consistently emerged as the most influential variable across all runs, with median weighted importance score of 0.35 (range: 0.21-0.43). Relative humidity was the second most important predictor, with median importance score of 0.24 (range: 0.16-0.33). Elevation, cropland, and climatic variables such as temperature seasonality (BIO04) and precipitation of the wettest quarter (BIO16) exhibited moderate importance, while other predictors showed low importance across all runs.

**Figure 2.**
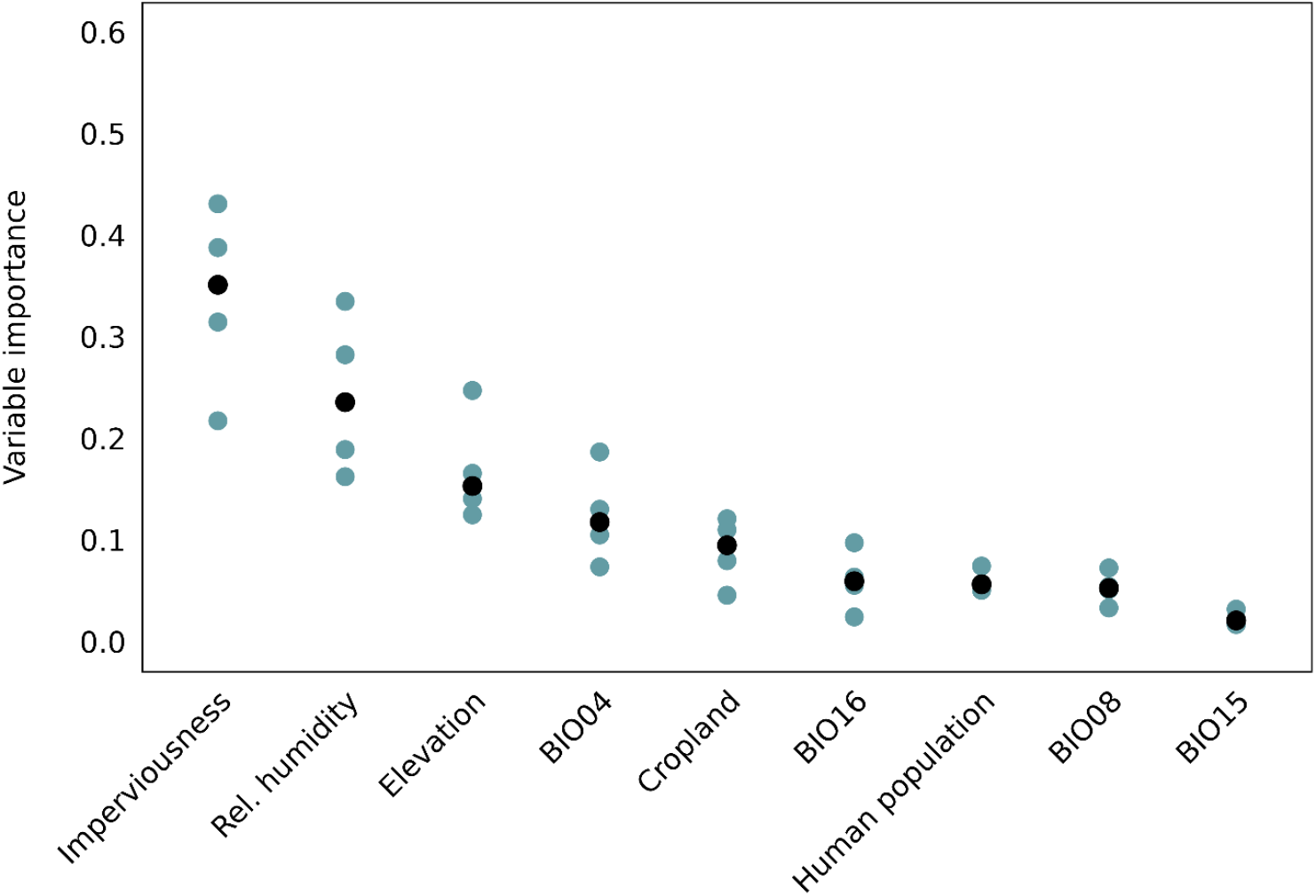
**Variable importance** values for each predictor (x-axis) calculated as the weighted mean of importance values for each run of the algorithms included in the ensemble (blue dots), and the median across all runs (black dots).

We plotted response curves to display the relationship between the predicted presence probability of *Cx. pipiens* and the covariates included in the model ensemble (Figure 3). The probability of presence sharply increased with imperviousness, highlighting a strong association with sealed surfaces. A similar positive response was observed with human population density, reinforcing the association between *Cx. pipiens* and anthropogenic environments. Relative humidity displayed a strong negative relationship with *Cx. pipiens* presence, with higher probability associated with drier conditions and a rapid decline as humidity increased. To further explore this relationship, we generated a bivariate response plot examining the interaction between relative humidity and precipitation of the wettest quarter (BIO16) in relation to predicted presence probability (Figure S4). The analysis revealed that under high precipitation conditions, humidity had little influence, with probabilities consistently exceeding 0.6. In contrast, at moderate to low precipitation levels, humidity became a key determinant, with presence probabilities decreasing below 0.3 in drier conditions.

**Figure 3.**
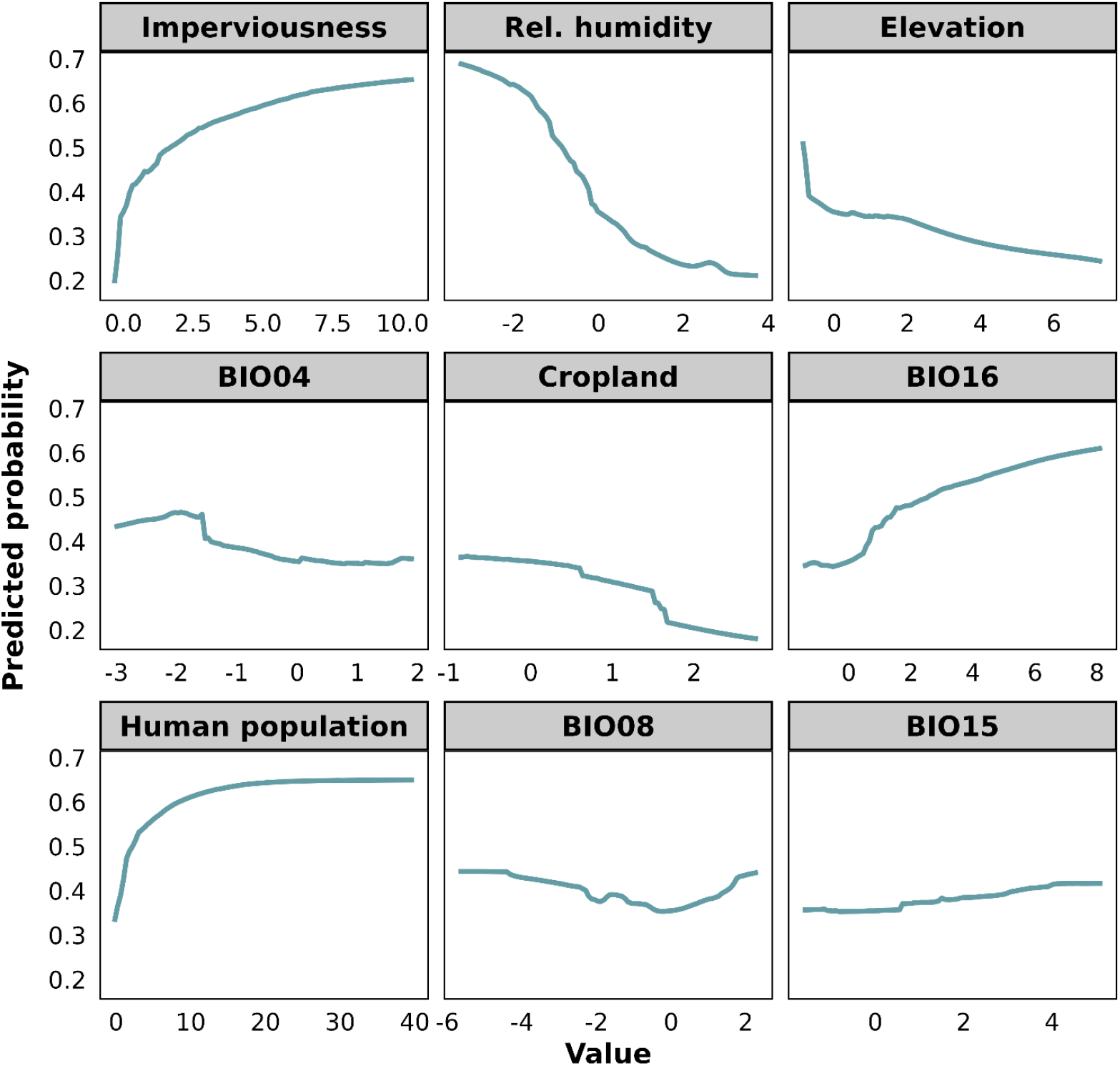
Response curves representing the relationship between the predicted presence probability of *Cx. pipiens* (on the y-axis) and different levels of explanatory variables (on the x-axis). For each curve, the effect of the variable of interest on the predicted distribution is isolated by fixing the other variables with the mean value.

The mean temperature of the wettest quarter (BIO08) followed a similar declining trend, reflecting a preference for cooler conditions during wet periods. Conversely, the positive effect of the precipitation of the wettest quarter (BIO16) suggested the affinity of *Cx. pipiens* for moderately wet environments. Elevation exhibited a marked negative relationship, underscoring the species’ preference for lowland habitats. Temperature seasonality (BIO04) showed a more complex, non-linear response, indicating idiosyncratic effects on the species’ distribution. Precipitation seasonality (BIO15) and cropland exhibited relatively flat curves.

### 3.3 Spatial predictions of habitat suitability

Based on the prediction of the ensemble model, we found several areas of high suitability for *Cx. pipiens* across Europe (Figure 4). High suitability regions are primarily distributed in heavily anthropized lowlands, reflecting the influence of imperviousness and elevation as key predictors. These areas can be found across all Europe, highlighting the widespread distribution potential of the species. When looking at the binarized map of species’ suitable habitat (Figure S5) we found that 25.6% of European terrestrial surface is potentially suitable for *Cx. pipiens*. Countries with the highest extent of suitable habitat for *Cx. pipiens* include Spain (238,276 km^2^), France (109,047 km^2^), and Italy (76,898 km^2^), and Germany (76,467 km^2^) (Table S4).

**Figure 4.**
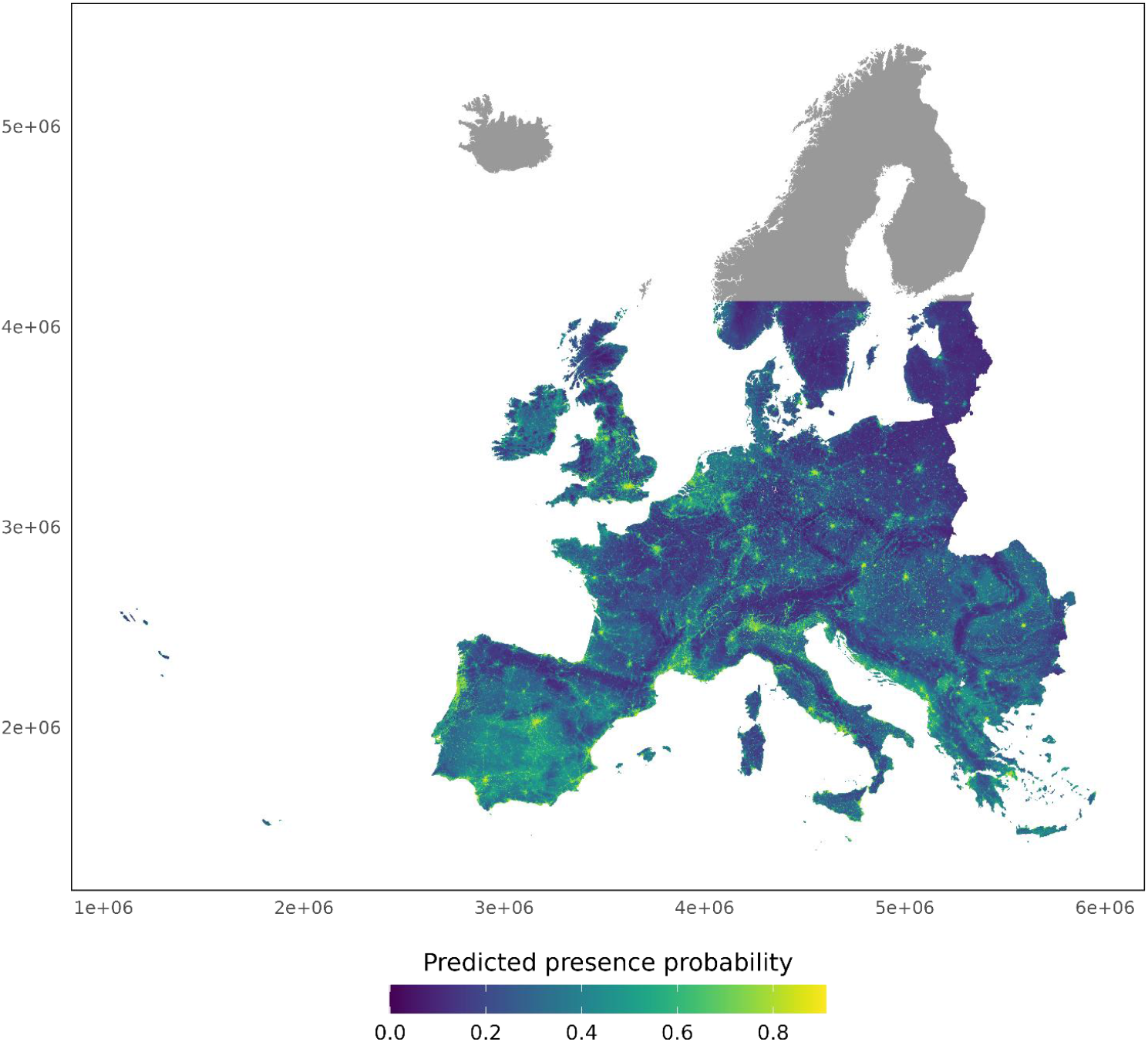
Distribution map of *Culex pipiens*. Predicted probability of presence of *Cx. pipiens* within the latitudinal limits encompassing 99% of *Cx. pipiens* occurrence records of the species.

## 4. Discussion

We implemented an ensemble modelling approach to explore the environmental factors shaping *Cx. pipiens* distribution across Europe. The ensemble approach outperformed individual models in predicting the habitat suitability of *Cx. pipiens*, addressing potential overfitting issues. Results from the new-label validation, conducted on independent data, further confirmed the strong predictive power of our model. Using this robust approach, we identified relevant anthropogenetic and climatic predictors contributing to the habitat suitability of *Cx. pipiens*, the primary vector for the transmission of several infections in humans, including West Nile virus. We found that habitat suitability for *Cx. pipiens* is widespread throughout Europe, without clear biogeographical pattern. *Cx. pipiens* distribution is primarily shaped by topography and the presence of human infrastructure, while bioclimatic variables played a more limited role.

An important finding from our study is the role of imperviousness in determining the distribution of *Cx. pipiens*. While several studies have identified urbanization as a contributing factor to the presence of *Cx. pipiens* at local scale (Gangoso et al., 2020; Rodríguez-Escolar et al., 2023), the explicit importance of imperviousness as a key predictor at continental scale emerged clearly from our analysis. The strong association between *Cx. pipiens* and human-modified areas underscores the need of considering anthropogenic factors in the modelling of vector species, particularly in highly modified environments. The tolerance of *Cx. pipiens* to temperature oscillations showed an idiosyncratic trend with a complex, non-linear effect of temperature seasonality (BIO04), and higher predicted presence probability for both high and low levels of temperature variability.

Imperviousness prevents rainwater from penetrating the soil, leading to increased surface runoff and a risk of stagnant water. At the same time, urbanization is associated with artificial water sources such as storm drains and water containers. These modifications of surface water regimes, combined with relatively stable thermal conditions in built-up areas, produce suitable microclimatic conditions for mosquito larvae and extend the breeding seasons (Becker et al., 2010). The relationship between elevation and *Cx. pipiens* presence highlights the species’ strong preference for lowland habitats. Lower elevations often coincide with higher human population densities and urbanized landscapes (Cohen & Small, 1998), where anthropogenic habitats provide a variety of opportunities for mosquito breeding and host-seeking behaviour (Becker et al., 2010).

Relative humidity exhibited a negative relationship with *Cx. pipiens* presence, indicating species’ preference for moderately dry environments. This might reflect its adaptability to urban microhabitats where water is available for breeding, but surrounding conditions are less humid compared to natural wetland ecosystems (Chakraborty et al., 2022). Urban areas often feature lower humidity due to alterations in land surfaces, such as increased impervious areas that facilitate rapid runoff and decreased vegetation cover, which limits evapotranspiration (Qian et al., 2022). Other factors might also contribute to the observed negative relationship between relative humidity and *Cx. pipiens* presence. In arid regions, human activities such as irrigation of gardens and green spaces may create localized breeding habitats, potentially leading to an interaction between humidity and population density (Ferraguti et al., 2016). Additionally, temperature could act as a confounding factor, as drier urban areas often experience higher temperatures (Yang et al., 2023), which may favour mosquito populations.

Compared to previous studies that employed species distribution models to map *Cx. pipiens* distribution in Europe (Versteirt et al., 2013; Wint et al., 2020), our results indicate a generally lower (albeit still substantial) amount of habitat suitability for *Cx. pipiens* across Europe. This difference may be attributed to our ensemble modelling framework, which integrates multiple algorithms, as well as our stricter validation approach and optimization techniques, which reduce overestimation of suitable areas. On the other hand, despite the more conservative nature of our predictions, showing a lower proportion of suitable areas for *Cx. pipiens* in Italy than those reported in previous works, our suitability map successfully identifies regions that are known to be suitable based on field studies and monitoring efforts, but had been misclassified as unsuitable, including areas of the Po Valley in Veneto and Emilia-Romagna (De Nardi et al., 2025; Mancini, 2017; Mulatti et al., 2014) where repeated outbreaks of West Nile virus have been recorded in the last decade (De Nardi et al., 2025).

Overall, this study underscores the importance of both anthropogenic and environmental factors in shaping the spatial distribution of *Cx. pipiens*. The strong association with urbanized areas and low-elevation zoned emphasizes the role of human-modified habitats in facilitating the spread of this vector species. Integrating these findings into surveillance and control strategies is essential for mitigating the risk of vector-borne diseases particularly in urban and periurban settings where *Cx. pipiens* thrives.

## 5. Acknowledgements

This research was supported by EU funding within the NextGeneration EU-MUR PNRR Extended Partnership initiative on emerging infectious diseases (project no. PE00000007, INF-ACT Spoke4).

## 6. Data availability

The maps produced in this work are available at https://zenodo.org/records/15040800.

