## Supplementary material for "Mapping the habitat suitability of *Culex pipiens* in Europe using ensemble bioclimatic modelling": Paper_Culex_SM.docx

*Lara Marcolin^1^*, Agnese Zardini^2^, Eleonora Longo^3^, Beniamino Caputo^3^, Piero Poletti^2^, Moreno Di Marco^1^*

^1^Department of Biology and Biotechnologies "Charles Darwin", Sapienza University of Rome, Italy

^2^Center for Health Emergencies, Bruno Kessler Foundation, Trento, Italy

^3^Department of Public Health and Infectious Diseases, Sapienza University of Rome, Italy.

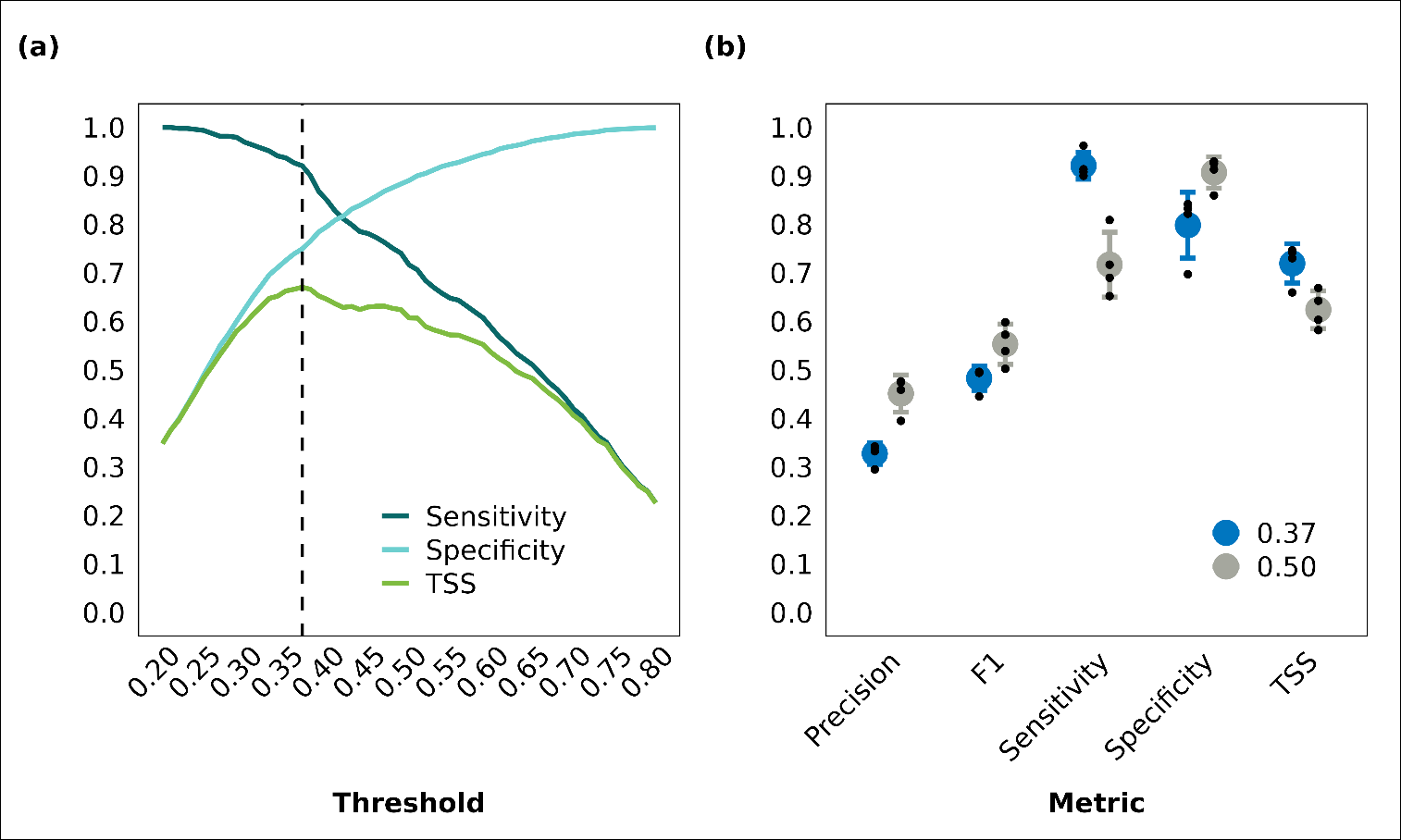

**Figure S1. Performance metrics of the model ensemble across different binarization thresholds.** (a) Sensitivity, specificity, and true skill statistic (TSS) values of the spatial block cross-validation averaged across the four blocks for different binarization thresholds. The dashed vertical line indicates the threshold (0.37) that maximizes the TSS. (b) Comparison of performance metrics (x-axis) of the ensemble model calculated using the default threshold (0.50) and the TSS-maximizing threshold (0.37). Performance metrics include precision (range: 0–1), F1 score (range: 0–1), sensitivity (range: 0–1), specificity (range: 0–1), and TSS (range: -1 to +1). Black dots correspond to spatial block cross-validation set runs.

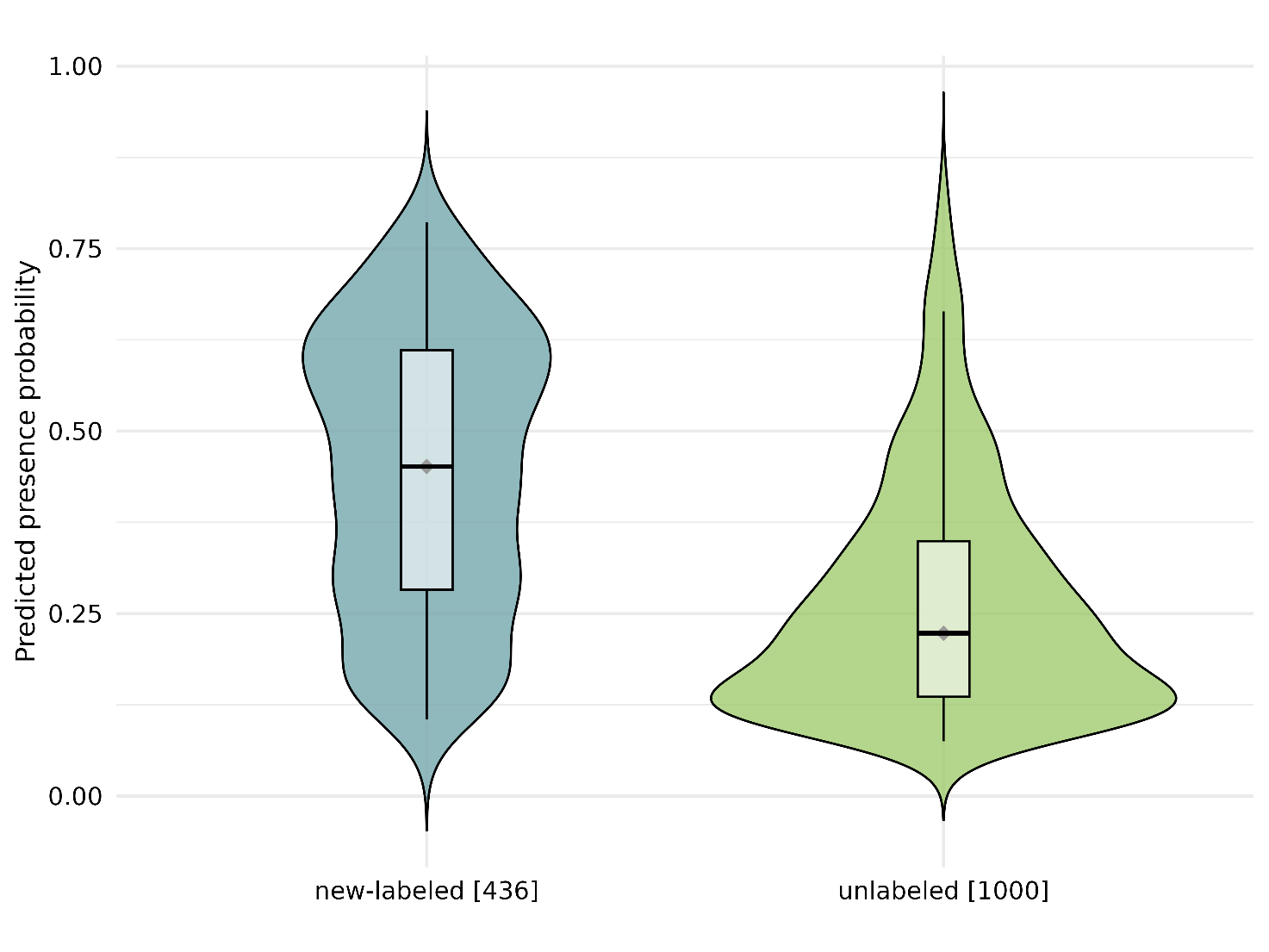

**Figure S2. Distribution of predicted presence probabilities** for occurrence points (“new-labeled” and background points (unlabeled). The violin plots show the probability of presence estimated for newly labeled occurrence points (n = 436) and unlabeled background points (n = 1000). The central boxplots indicate the median and interquartile range. The newly labeled occurrences exhibit a higher median probability of presence compared to background points.

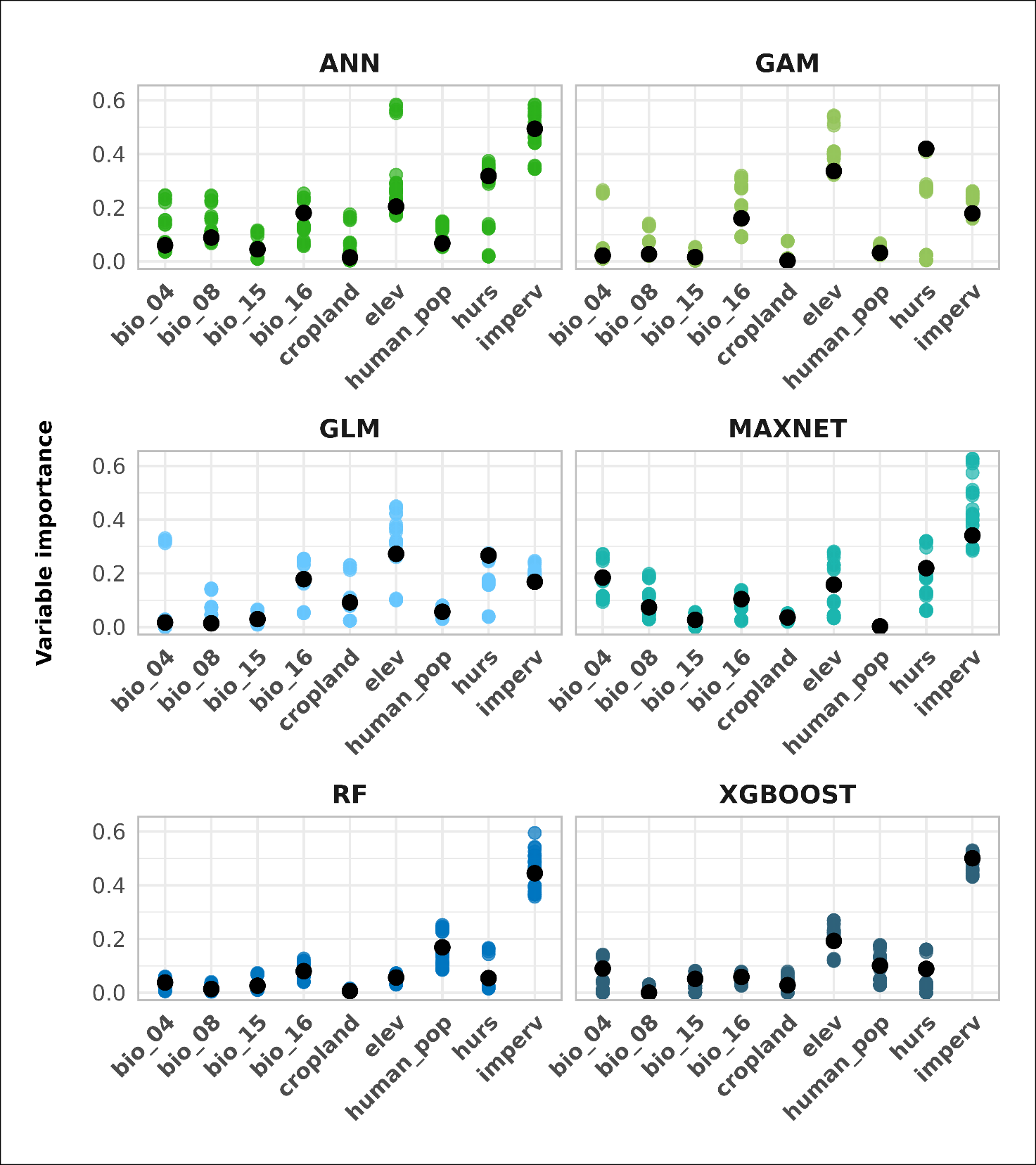

**Figure S3.** **Variable importance values** for each predictor (x-axis) calculated using permutation importance with five permutations for each run of the algorithms included in the ensemble model. Each dot represents a single permutation result, with different panels corresponding to different algorithms. Black dots represent the mean importance value for each predictor variable across all runs and permutations.

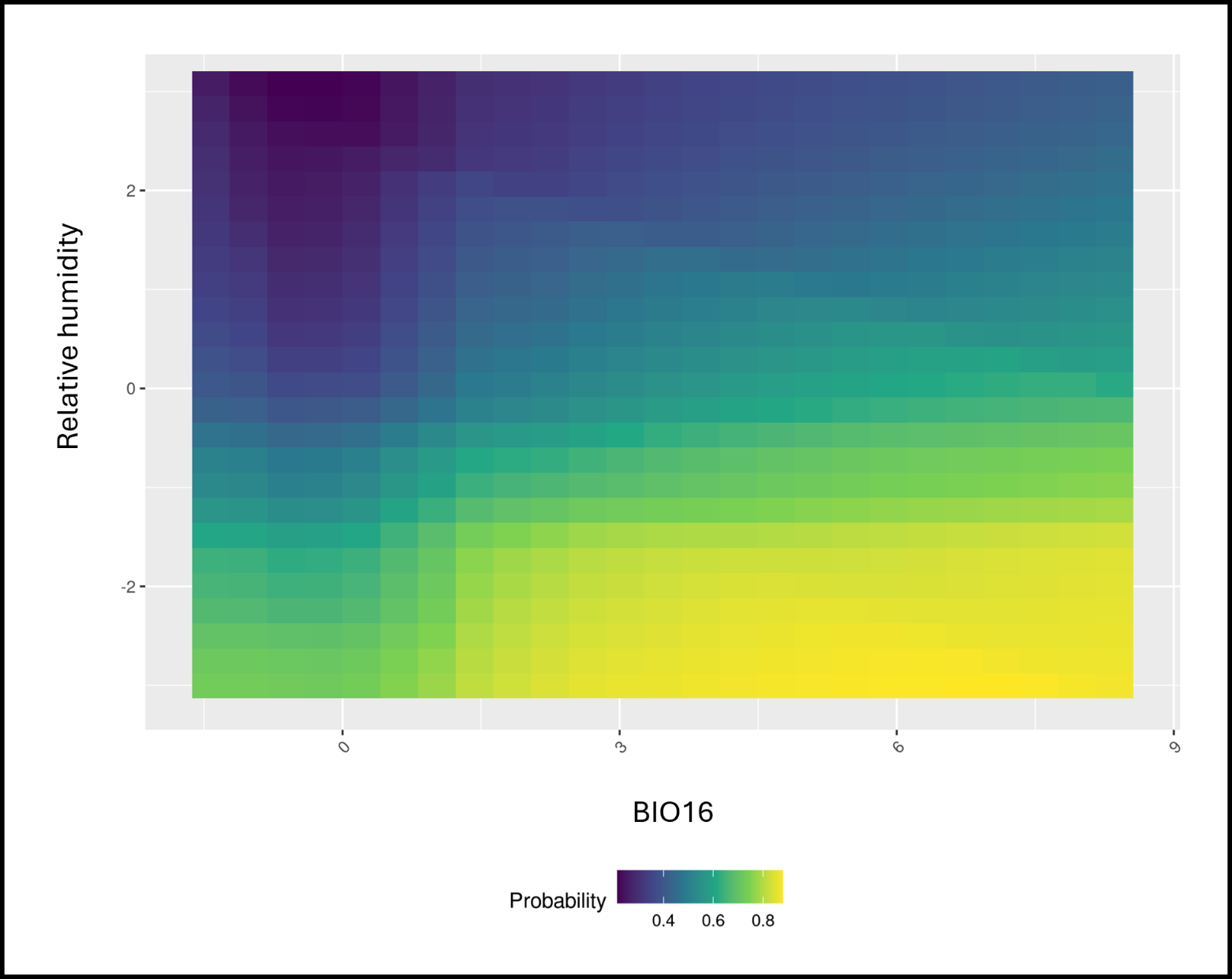

**Figure S4. Bivariate response plot** showing the effect of the interaction of relative humidity and precipitation of the wettest quarter (BIO16) on the predicted probability of *C. pipiens* presence.

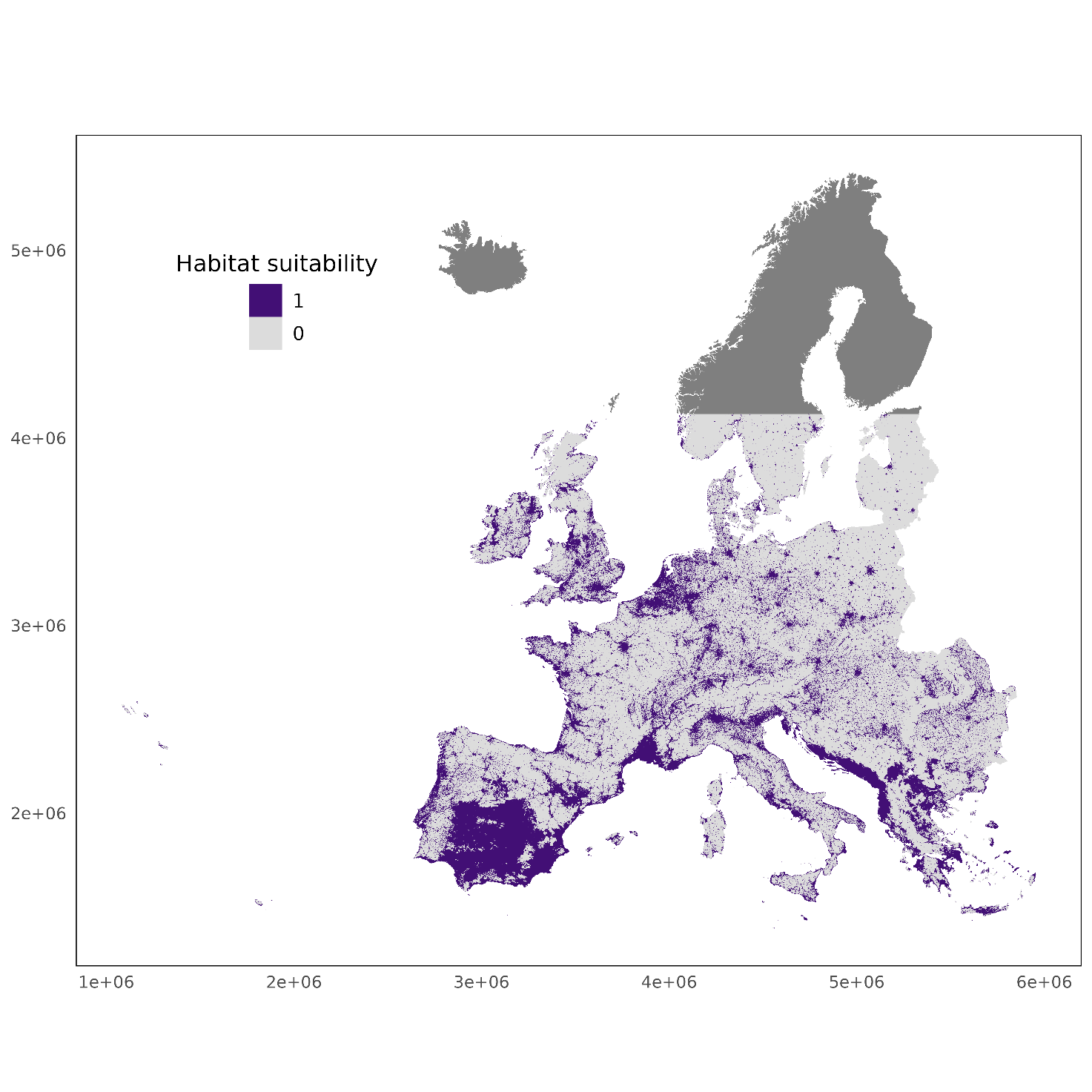

**Figure S5. Predicted habitat suitability map** of *Culex pipiens* obtained with a binarization threshold of 0.37 within the latitudinal limits encompassing 99% of *C. pipiens* occurrence records of the species.

**Table S1. Algorithms included in the model ensemble** and corresponding parameter settings.

| **Algorithm** | **Hyperparameter** | **Value** | **Description (Thuiller et al., 2016)** |
| --- | --- | --- | --- |
| Random Forest | mtry  min_n | 5  5 | Number of features that will be randomly sampled at each split when creating the tree models  Minimum number of data points in a node that are required for the node to be split further |
| Extreme gradient boosting | max_depth  eta | 6  0.3 | Maximum depth of a tree  Step size shrinkage used in update to prevent overfitting |
| Generalized additive model | k | 3 | The number of basis functions used for a smooth term. |
| Neural Network | size  decay | 2  1e-03 | Number of units in the hidden layer  Parameter for weight decay |
| Maxnet | regmult | 3 | A constant to adjust regularization |

**Table S2. Variance inflation factor** (VIF) of explanatory variables included in the models. VIF is derived from the square of the multiple correlation coefficient obtained by regressing a predictor on all other predictors. When a variable shows a strong linear relationship with one or more variables, the correlation coefficient is close to 1, leading to a high VIF for that predictor.

| **Explanatory variable** | **Variance inflation factor** |
| --- | --- |
| BIO04 | 1.491 |
| BIO08 | 2.070 |
| BIO15 | 1.757 |
| BIO16 | 1.841 |
| Elevation | 1.954 |
| Relative humidity | 1.909 |
| Imperviousness | 1.639 |
| Human population | 1.545 |
| Cropland | 1.387 |

**Table S3. Model evaluation.** Boyce index and TSS scores for all runs included in the ensemble model.

| **Run** | **Algorithm** | **Metric** | **Cutoff** | **Calibration** | **Validation** |
| --- | --- | --- | --- | --- | --- |
| RUN1 | RF | BOYCE | 0.399 | 0.93 | 0.96 |
| RUN1 | RF | TSS | 0.334 | 0.99 | 0.63 |
| RUN1 | MAXNET | BOYCE | 0.327 | 0.99 | 0.98 |
| RUN1 | MAXNET | TSS | 0.334 | 0.65 | 0.45 |
| RUN1 | XGBOOST | BOYCE | 0.494 | 0.99 | 0.82 |
| RUN1 | XGBOOST | TSS | 0.508 | 0.59 | 0.52 |
| RUN1 | ANN | BOYCE | 0.400 | 0.97 | 0.67 |
| RUN1 | ANN | TSS | 0.430 | 0.59 | 0.39 |
| RUN1 | GAM | BOYCE | 0.455 | 0.99 | 0.70 |
| RUN1 | GAM | TSS | 0.433 | 0.47 | 0.10 |
| RUN2 | RF | BOYCE | 0.339 | 1.00 | 0.81 |
| RUN2 | RF | TSS | 0.318 | 0.99 | 0.37 |
| RUN2 | MAXNET | BOYCE | 0.307 | 0.99 | 0.90 |
| RUN2 | MAXNET | TSS | 0.321 | 0.66 | 0.45 |
| RUN2 | XGBOOST | BOYCE | 0.498 | 0.97 | 0.81 |
| RUN2 | XGBOOST | TSS | 0.343 | 0.60 | 0.29 |
| RUN2 | ANN | BOYCE | 0.475 | 0.93 | 0.83 |
| RUN2 | ANN | TSS | 0.436 | 0.59 | 0.40 |
| RUN2 | GAM | BOYCE | 0.446 | 0.99 | 0.92 |
| RUN2 | GAM | TSS | 0.414 | 0.45 | 0.27 |
| RUN3 | RF | BOYCE | 0.331 | 0.99 | 0.93 |
| RUN3 | RF | TSS | 0.346 | 0.99 | 0.29 |
| RUN3 | MAXNET | BOYCE | 0.280 | 0.99 | 0.95 |
| RUN3 | MAXNET | TSS | 0.264 | 0.64 | 0.41 |
| RUN3 | XGBOOST | BOYCE | 0.467 | 0.97 | 0.70 |
| RUN3 | XGBOOST | TSS | 0.439 | 0.62 | 0.34 |
| RUN3 | ANN | BOYCE | 0.657 | 0.77 | 0.80 |
| RUN3 | ANN | TSS | 0.640 | 0.57 | 0.34 |
| RUN3 | GAM | BOYCE | 0.524 | 0.99 | 0.80 |
| RUN3 | GAM | TSS | 0.491 | 0.51 | 0.35 |
| RUN4 | RF | BOYCE | 0.389 | 0.99 | 0.94 |
| RUN4 | RF | TSS | 0.407 | 0.99 | 0.29 |
| RUN4 | MAXNET | BOYCE | 0.363 | 0.99 | 0.99 |
| RUN4 | MAXNET | TSS | 0.366 | 0.59 | 0.39 |
| RUN4 | XGBOOST | BOYCE | 0.646 | 0.96 | 0.86 |
| RUN4 | XGBOOST | TSS | 0.577 | 0.62 | 0.37 |
| RUN4 | ANN | BOYCE | 0.496 | 0.97 | 0.71 |
| RUN4 | ANN | TSS | 0.500 | 0.59 | 0.29 |
| RUN4 | GAM | BOYCE | 0.611 | 0.97 | 0.86 |
| RUN4 | GAM | TSS | 0.625 | 0.45 | 0.22 |

**Table S4. Proportion and total area** of predicted habitat suitability for each European country.

| **Country** | **Proportion (%)** | **Area (km^2^)** |
| --- | --- | --- |
| Albania | 54.20 | 13863 |
| Andorra | 8.94 | 39 |
| Austria | 16.43 | 12536 |
| Belgium | 42.49 | 11863 |
| Bosnia and Herz. | 20.17 | 9475 |
| Bulgaria | 15.98 | 16213 |
| Croatia | 34.38 | 17181 |
| Czechia | 13.61 | 9701 |
| Denmark | 20.60 | 7678 |
| Estonia | 1.86 | 756 |
| France | 22.27 | 109047 |
| Germany | 23.88 | 76467 |
| Greece | 43.49 | 50863 |
| Hungary | 15.59 | 13116 |
| Ireland | 28.99 | 17984 |
| Italy | 28.55 | 76898 |
| Kosovo | 27.98 | 5625 |
| Latvia | 2.32 | 1218 |
| Liechtenstein | 53.74 | 79 |
| Lithuania | 2.84 | 1653 |
| Luxembourg | 18.72 | 463 |
| Malta | 92.05 | 255 |
| Moldova | 51.27 | 322 |
| Monaco | 88.23 | 15 |
| Montenegro | 47.31 | 5935 |
| Netherlands | 61.88 | 19771 |
| North Macedonia | 45.16 | 10420 |
| Norway | 3.82 | 10871 |
| Poland | 8.53 | 23934 |
| Portugal | 36.50 | 29741 |
| Romania | 16.33 | 34640 |
| San Marino | 70.13 | 54 |
| Serbia | 15.95 | 11163 |
| Slovakia | 10.83 | 4763 |
| Slovenia | 25.36 | 4705 |
| Spain | 53.36 | 238276 |
| Sweden | 1.86 | 7442 |
| Switzerland | 23.17 | 8732 |
| United Kingdom | 28.59 | 61680 |
| Vatican | 100 | 1 |
